# Temporal processing of facial expressions of mental states

**DOI:** 10.1101/602375

**Authors:** Gunnar Schmidtmann, Joshua T. Loong, Claus-Christian Carbon, Maiya Jordan, Andrew J. Logan, Ian Gold

## Abstract

Faces provide not only cues to an individual’s identity, age, gender and ethnicity, but also insight into their mental states. The ability to identify the mental states of others is known as Theory of Mind. Here we present results from a study aimed at extending our understanding of differences in the temporal dynamics of the recognition of expressions beyond the basic emotions at short presentation times ranging from 12.5 to 100 ms. We measured the effect of variations in presentation time on identification accuracy for 36 different facial expressions of mental states based on the Reading the Mind in the Eyes test (Baron-Cohen et al., 2001) and compared these results to those for corresponding stimuli from the McGill Face database, a new set of images depicting mental states portrayed by professional actors. Our results show that subjects are able to identify facial expressions of complex mental states at very brief presentation times. The kind of cognition involved in the correct identification of facial expressions of complex mental states at very short presentation times suggests a fast, automatic Type-1 cognition.

## Introduction

Faces are amongst the most complex objects processed by the human visual system and contain a wealth of information (Bruce & Young, 1986). In addition to providing cues to an individual’s identity, age, gender and ethnicity, faces provide insight into an individual’s emotions, beliefs and intentions. In other words, faces are the central focus for our ability to attribute mental states to others - that is, for our Theory of Mind capacity. As such, facial expressions are an important component of non-verbal communication and provide information which is critical for social interactions (Ekman, 1973). In processing facial expressions, humans recruit an expansive network of brain regions, including the superior temporal sulcus, orbitofrontal cortex, insular cortex and the amygdala (Haxby et al., 2000, Phillips et al., 1997, Pitcher et al., 2017). A rich Theory of Mind capacity appears to be unique to humans (Devaine et al., 2017) and is thought to be essential to human social behaviour. Impaired Theory of Mind capacity is associated with a variety of clinical conditions, notably autism (Baron-Cohen, 1997, Baron-Cohen et al., 1985, 2001) and schizophrenia (Frith, 2014).

Facial expressions engage rapid visual processing mechanisms; less than 100 ms of exposure is typically sufficient to identify facial expressions of emotion (Calvo & Esteves, 2005, Neath & Itier, 2014). A prevailing view is that, in response to evolutionary needs, humans have developed the ability to express and identify a small number of basic emotions; happiness, sadness, fear, surprise, disgust, anger and contempt (Ekman, 1992, Ekman & Cordaro, 2011). This is supported by behavioural evidence that these expressions are reliably recognized across a range of stimuli and paradigms as well as across cultures (Calvo & Lundqvist, 2008, Elfenbein & Ambady, 2002, Palermo & Coltheart, 2004), and neuroimaging studies which have identified dissociable neural correlates of these basic emotions (Adolphs, 2002, Hamann, 2012, Vytal & Hamann, 2010).

A wealth of evidence, however, points to the view that there are significant differences in the time-course of processing different facial expressions (e.g. happiness and anger) (Milders et al., 2008, Nummenmaa & Calvo, 2015, Palermo & Coltheart, 2004). For example, a number of studies have utilized a simultaneous discrimination paradigm to demonstrate that angry faces are detected more rapidly than either happy or neutral faces (Fox et al., 2000, Hansen & Hansen, 1988, Öhman et al., 2001, Sato & Yoshikawa, 2010, Sawada et al., 2014). In line with this behavioural evidence, it has been reported that the latencies of ERP components associated with facial expression processing are significantly shorter for angry relative to happy faces (Feldmann-Wüstefeld et al., 2011). It has been proposed that these differences in perceptual latency reflect the ecological salience of particular emotions (Li et al., 2018). Specifically, rapid detection of a facial expression which communicates threat (e.g. an angry face) may be advantageous for survival (Öhman et al., 2001). It should be noted, however, that a number of studies have found no evidence of a detection advantage for angry faces (Becker et al., 2011, Calvo & Nummenmaa, 2008). It has been suggested that these conflicting results can be explained by methodological differences across several studies (Nummenmaa & Calvo, 2015), where detection advantage is rather determined by the perceptual features of an image (for example, contrast).

The detection of facial expressions is a cognitive process distinct from expression identification, and identification involves further processing beyond that required for detection. The angry-advantage for the detection of emotion notwithstanding, a good deal of evidence points to the conclusion that positive facial expressions (e.g. happiness) are identified more readily than negative expressions (e.g. sadness) (Calvo & Lundqvist, 2008, Leppänen & Hietanen, 2004, Milders et al., 2008, Neath & Itier, 2014, Palermo & Coltheart, 2004). For example, Calvo & Lundqvist (2008) found that happy expressions were identified more accurately and rapidly than faces exhibiting sadness, anger or fear. This happiness-advantage persists when schematic faces are utilized to control for physical differences between happy and sad faces Leppänen & Hietanen (2004). It has been proposed that the happiness-advantage can be explained by the fact that happiness is the only pleasant basic emotion or by the upturned mouth that is unique to its facial manifestation (Nummenmaa & Calvo, 2015). At the other end of the spectrum, it has been reported that fear is recognized more slowly than the other basic emotions (Calvo et al., 2014, Calvo & Lundqvist, 2008, Palermo & Coltheart, 2004, Tracy & Robins, 2008). The evidence for this fear-disadvantage, however, is mixed; other studies have found no difference between the processing of fearful, angry, neutral and sad facial expressions (Calvo & Nummenmaa, 2009, Milders et al., 2008).

It has also been proposed that, rather than being categorized by means of discrete categories such as happiness or sadness, facial expressions are better described with respect to continuous variables along different dimensions, e.g. arousal and valence (Russell, 1994, Takehara & Suzuki, 1997). For example, an angry face is high in arousal but low in valence, while a bored face is low in arousal and average in valence. In support of this latter view in particular, participants’ ratings of facial expressions have been found to be successfully captured by two independent dimensions, valence and arousal (Takehara & Suzuki, 1997). Further, describing facial expressions in terms of dimensions, rather than discrete categories, is more consistent with evidence of differences in perceived facial expression intensity (Hess et al., 1997). More recently, it has been proposed that humans use both categorial and dimensional approaches to process facial expressions (Fujimura et al., 2012, Harris et al., 2012).

Nevertheless, most previous investigations of the temporal properties of facial expression processing have focused upon a small number of basic emotions. The Theory of Mind capacity is directed not only at those states but at the social emotions (e.g., shame), as well as non-emotional states such as desire, belief, intention, and the like. One would do well to be cautious about generalizing models of the perception of the basic emotions to these other facial expressions. In particular, the temporal dynamics of the perception of facial expressions beyond the basic emotions remains an open question. What those dynamics are will have implications for modelling the Theory of Mind capacity.

Over the last nearly 40 years, a number of models of Theory of Mind have been developed and the mainstream views are similar in that they tend to construe Theory of Mind as a form of cognition that is demanding and slow. For instance, “Theory-Theory” understands Theory of Mind as supported by a process of reasoning about others by means of the deployment of a folk psychological theory (for a historical review of the approaches to Theory of Mind, see Goldman et al. (2012)). On the analogy of scientific reasoning, the mental states of others are inferred on the basis of law-like psychological generalizations together with the particular conditions in which the agent finds herself. In contrast, “Simulation-Theory” posits that Theory of Mind is exercised by means of an imaginative identification with the person whose mental states are of interest. By putting oneself in the shoes of another person, one can take on their mental states imaginatively as one’s own. Each of these models construe Theory of Mind as being computationally resource-intensive and predicts that Theory of Mind should typically be slow (Apperly, 2010).

Recently, however, a number of investigators have noted that the everyday use of Theory of Mind has to happen “on the fly” - automatically, effortlessly and quickly - for social interaction to proceed unimpeded (e.g., Gallagher, 2001). A proposal by Apperly and Butterfly (2009) offers an explanation of this apparent conflict between theory and observation. According to their hypothesis, Theory of Mind capacity is subserved by two distinct processes - one that is (among other things) automatic, effortless, and fast, and a second that is consciously engaged, effortful, and slow. The idea that a single cognitive goal is subserved by two systems - one that is “designed” to provide immediate, rough-and-ready solutions and another one that is slower but more systematic - is known as “dual-process” or “dual-system” theories, with “Type 1” cognition referring to the function of the former system or process, and “Type 2” cognition referring to the latter (Evans, 2008).

The present study aimed to elucidate differences in the temporal aspects of the processing of a wide range of facial expressions. Since speed is a symptom of Type 1 cognition, if Theory of Mind can be shown to happen quickly, this would provide some evidence for the merit of a dual-process understanding of Theory of Mind. At the very least, evidence of very fast Theory of Mind capacity seems to call out for some explanation beyond Theory-Theory or Simulation-Theory. As already noted, the majority of previous studies have investigated the time-course of facial expression processing for only a limited number of basic emotions (e.g. happiness, sadness, fear, surprise, disgust and anger). To our knowledge, the ability to identify nuanced differences between facial expressions which are considerably more similar, in terms of arousal and valence ratings, than the basic emotional categories (e.g. happiness or anger) has not been tested. For example, it has not been established whether humans can reliably identify differences between facial expressions of upset, despondence and disappointment. Further, the temporal aspects of processing these complex mental states have not been investigated. The present study aimed to extend understanding of differences in the time-course of processing facial expressions beyond the basic emotions at fast presentation times.

## Methods

### Subjects

All participants were recruited through a McGill Facebook group. Thirty individuals (20 females, 2 non-binary, 8 males, mean age 21 years, range: 18 − 30 years) participated in the study. All participant reported at least ten years of English fluency and were naïve as to the purpose of the study. Participants reported normal, or corrected-to-normal, vision. Written informed consent was obtained from each participant. Further details are summarized in Table 1. All experiments were approved by the McGill University Ethics committee (dossier number 51-0714) and were conducted in accordance with the original Declaration of Helsinki.

**Table 1.** Subject details

| **Subject ID** | **Age** | **Gender** | **Race** | **Religion** | **Country of Origin** | **Years in Canada** | **Guess Rate** |
| --- | --- | --- | --- | --- | --- | --- | --- |
| 1 | 23 | Male | Asian | Hinduism | India | 11 | 60 – 70 (65) |
| 2 | 22 | Female | White | None | Canada | 22 | 40 |
| 3 | 21 | Female | Multiracial | None | Brazil | 0.83 | 60 |
| 4 | 24 | Female | Other | None | Iran | 22 | 40 |
| 5 | 19 | Female | White/Asian | Christianity | Canada | 19 | 50 |
| 6 | 30 | Male | White | Christianity | Canada | 30 | 40 |
| 7 | 23 | Female | White | None | Canada | 23 | 75 |
| 8 | 24 | Female | White | Christianity | Canada | 24 | 50 |
| 9 | 20 | Male | White | None | Canada | 20 | 65 |
| 10 | 20 | Female | Asian | None | China | 14 | 40 |
| 11 | 19 | Female | White | Christianity | France | 3 | 3 |
| 12 | 19 | Female | Hispanic | None | Colombia | 16 | 90 |
| 13 | 20 | Male | White/Hispanic | None | Multicultural | 8 | 55 |
| 14 | 18 | Female | Asian | None | Vietnam | 1 | 55 |
| 15 | 21 | Male | Asian | None | Vietnam | 17 | 75 |
| 16 | 19 | Male | Asian | None | China | 1 | 50 |
| 17 | 24 | Female | Asian | None | China | 11 | 60 |
| 18 | 23 | Female | Hispanic | Other | Peru | 4 | 50 |
| 19 | 23 | Male | Other | Christianity | Peru | 3 | 60 |
| 20 | 22 | Female | Asian | Buddhism | China | 17 | 30 |
| 21 | 22 | Female | Asian | Christianity | Korea | 22 | 65 |
| 22 | 21 | Female | White | None | Russia | 9 | 50 |
| 23 | 20 | Male | White | None | Canada | 20 | 40 |
| 24 | 21 | Female | Asian | None | Canada | 21 | 35 |
| 25 | 20 | Female | Black | Christianity | Canada | 20 | 55 |
| 26 | 22 | Non-binary | White | None | United States | 25 | 60 |
| 27 | 21 | Non-binary | White/Asian | None | United States | 3 | 75 |
| 28 | 19 | Female | Black | Christianity | United States | 4 | 35 – 40 (37.5) |
| 29 | 20 | Female | White | Judaism | United States | 2 | 15 |
| 30 | 20 | Female | White | None | Canada | 20 | 25 |

### Apparatus

Experiments were performed in a dimly illuminated room. Stimuli were presented, using MATLAB (MATLAB R 2018b, MathWorks) and routines from the Psychtoolbox-3 (Brainard, 1997), on a gamma-corrected Mitsubishi DiamondPro 2070 CRT monitor with a resolution of 1280 × 1024 pixels and a frame rate of 80Hz (mean luminance: 60 *cd*/*m*^2^). The monitor was controlled by a MacBook Air computer (2015, 1.6 GHz). Participants viewed the stimuli at a distance of 55 cm. At this distance, one pixel subtended 0.037° visual angle.

### Stimuli

We employed a digital version of the original RMET stimuli (Baron-Cohen et al., 2001). The original RMET stimuli consisted of 36 black-and-white photographs of the eye region of both male and female individuals. The original stimuli were created by extracting photographs from magazines and were therefore not controlled for variables such as luminance, brightness, scale and perspective. To create a digital version of this test, we acuired electronic copies of the original photgraphs. The resultant stimuli were presented in the center of a mid-grey backgroud. The images were JPEG files with a resolution of 1024 × 768 pixel. When viewed at the test distance, the stimuli had a size of 10.4° × 4.0° of visual angle. The McGill Face stimuli were taken from the McGill Face Database (Schmidtmann et al., 2019). The full McGill Face Database contains pictures of 93 different expressions of mental states portrayed by two (one male, one female) English-speaking professional actors. For this study, we extracted 36 images from the full McGill Face Database which match the expressions portrayed in the RMET stimuli. The McGill stimuli were cropped to isolate the eye region and adjusted to match the size of the RMET stimuli. The full set of stimuli used in this study can be downloaded here: *Stimuli* The full McGill Face database can be downloaded at: *McGill Face Database*

The images used in the RMET were specifically chosen for inclusion within a test of sensitivity to the eye region. The McGill stimuli, on the other hand, were captured for a full-face test of facial expression sensitivity, without any specific focus being placed on the ocular region.

### Terms

All 36 target terms from the RMET were tested (Baron-Cohen et al., 2001). The three alternative terms for each target were those established and utilized by Baron-Cohen et al. (2001) (shown in their Appendix A). A list of the target terms with the corresponding alternative terms is shown in Table 2. Note that the terms cautious, fantasizing, interested, and preoccupied occur twice in the RMET (referred to as (a) and (b)).

**Table 2.** List of target and alternative terms taken from Baron-Cohen et al. (2001).

|  | **Target Term** | **Alternative 1** | **Alternative 2** | **Alternative 3** |
| --- | --- | --- | --- | --- |
| 1 | playful | comforting | irritated | bored |
| 2 | upset | terrified | arrogant | annoyed |
| 3 | desire | joking | flustered | convinced |
| 4 | insisting | joking | amused | relaxed |
| 5 | worried | irritated | sarcastic | friendly |
| 6 | fantasizing (a) | aghast | impatient | alarmed |
| 7 | uneasy | apologetic | friendly | dispirited |
| 8 | despondent | relieved | shy | excited |
| 9 | preoccupied (a) | annoyed | hostile | horrified |
| 10 | cautious (a) | insisting | bored | aghast |
| 11 | regretful | terrified | amused | flirtatious |
| 12 | sceptical | indifferent | embarrassed | dispirited |
| 13 | anticipating | decisive | threatening | shy |
| 14 | accusing | irritated | disappointed | depressed |
| 15 | contemplative | flustered | encouraging | amused |
| 16 | thoughtful | irritated | encouraging | sympathetic |
| 17 | doubtful | affectionate | playful | aghast |
| 18 | decisive | amused | aghast | bored |
| 19 | tentative | arrogant | grateful | sarcastic |
| 20 | friendly | dominant | guilty | horrified |
| 21 | fantasizing (b) | embarrassed | confused | panicked |
| 22 | preoccupied (b) | grateful | insisting | imploring |
| 23 | defiant | contended | apologetic | curious |
| 24 | pensive | irritated | excited | hostile |
| 25 | interested (a) | panicked | incredulous | despondent |
| 26 | hostile | alarmed | shy | anxious |
| 27 | cautious (b) | joking | arrogant | reassuring |
| 28 | interested (b) | joking | affectionate | contended |
| 29 | reflective | impatient | aghast | irritated |
| 30 | flirtatious | grateful | hostile | disappointed |
| 31 | confident | ashamed | joking | dispirited |
| 32 | serious | ashamed | bewildered | alarmed |
| 33 | concerned | embarrassed | guilty | fantasizing |
| 34 | distrustful | aghast | baffled | terrified |
| 35 | nervous | puzzled | insisting | contemplative |
| 36 | suspicious | ashamed | nervous | indecisive |

### Procedure

Participants were initially familiarized with each of the 93 terms which appeared within the test and their corresponding descriptions. Descriptions were extracted from the Glossary of Appendix B in Baron-Cohen et al. (2001). We employed the same four-alternative forced choice paradigm as that utilized within the original RMET task (Baron-Cohen et al., 2001). The experimental block began with presentation of a mid-grey background. On each trial, participants were shown the eye region of a face portraying an expression for one of eight specific presentation times (ms). Within a block, each expression was tested twice at each presentation time (2 × 8 × 36 = 576 trials per block). Expressions and presentation times were presented in a random order using an interleaved design. The RMET and McGill stimuli were tested in separate blocks. The stimulus was followed immediately by a mask (presented for 500ms, 8.8° × 8.8° of visual angle), which comprised random greyscale luminance noise. The purpose of the mask was to remove any residual visual transient. Following offset of the mask, participants were presented with four terms (font type: Arial, font color: white, font size: 60), arranged in a diamond format, on the midgrey screen. One of the terms (target) described the expression being portrayed by the stimulus. The remaining three terms (distractors) were those established by Baron-Cohen et al. (2001) (Alternatives terms in Table 2). The position of the target term (up, down, right or left) within the diamond configuration was randomly determined on each trial. The participant was asked to choose the term which best described the expression being portrayed by the stimulus. Response was indicated via keyboard press. No feedback was provided. After an experimental block each observer was asked to estimate the proportion of trials on which they felt that they were guessing the correct term. The guess rates are shown in the rightmost column of Table 1.

## Results

Figure 1 shows performance (response accuracy in proportion correct) as a function of presentation time ranging from 12.5 to 100 ms for the McGill stimuli (Figure 1A) and the RMET stimuli (Figure 1B). The small circular data points show the average individual performance for each subject and the blue solid line represents the average performance presented as a higher order polynomial regression model fit to the data.

**Figure 1.**
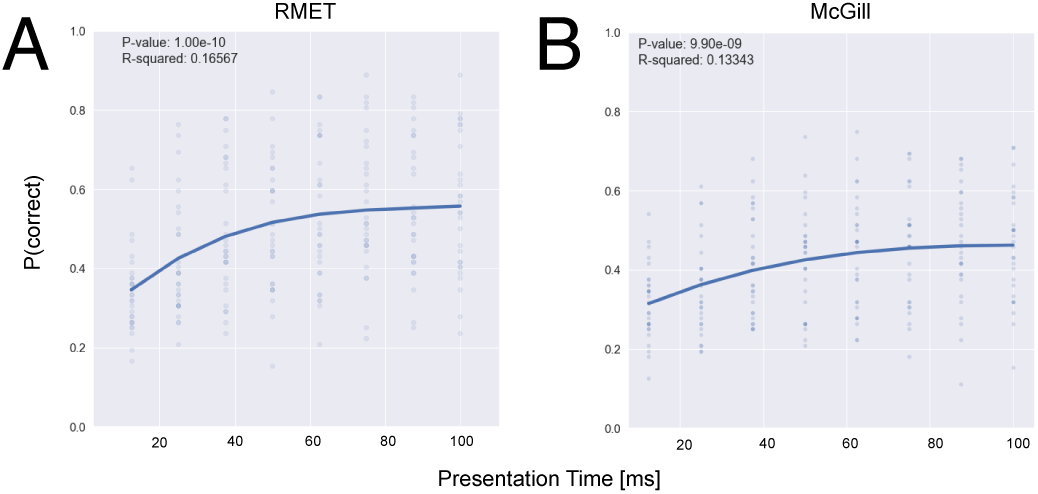
The figure shows performance (proportion correct) as a function of presentation time for (A) RMET and (B) the McGill Database stimuli. The small circular data points represent mean individual data and the solid blue line shows a power function fit to the data.

Results show that the average performance increases with increasing presentation time for both stimulus classes (RMET and McGill). In both conditions, a third-degree polynomial function was fit to the data (coefficient of determination *R*^2^and*p*-values are shown in each graph in Figure 1). Analysis of covariance of these polynomial regressions did not reveal statistically significant differences in performance between the two stimuli classes. This was in respect to both the intercept (*F*_2,692_ = 0.2063, *p* =. 650) and the slope parameters (*F*_2,692_ = 2.890, *p* =. 090). *χ*^2^ - tests with a Yates correction for continuity (*p* >. 05) revealed that on average, participants performed better than chance across all presentation times for both types of stimuli, and this effect becomes more pronounced as presentation time increases. Performance increased to over 70% of participants for longer presentation times. A one-way ANCOVA between the two stimuli classes revealed statistically significant differences in performance predicated by the model at the y-intercept better than chance performance (*F*_2,13_ = 59.942, *p* <. 001). However, the regression slopes shown in Figure 2 are not statistically significant (*F*_2,13_ = 0.045, *p* =. 834).

**Figure 2.**
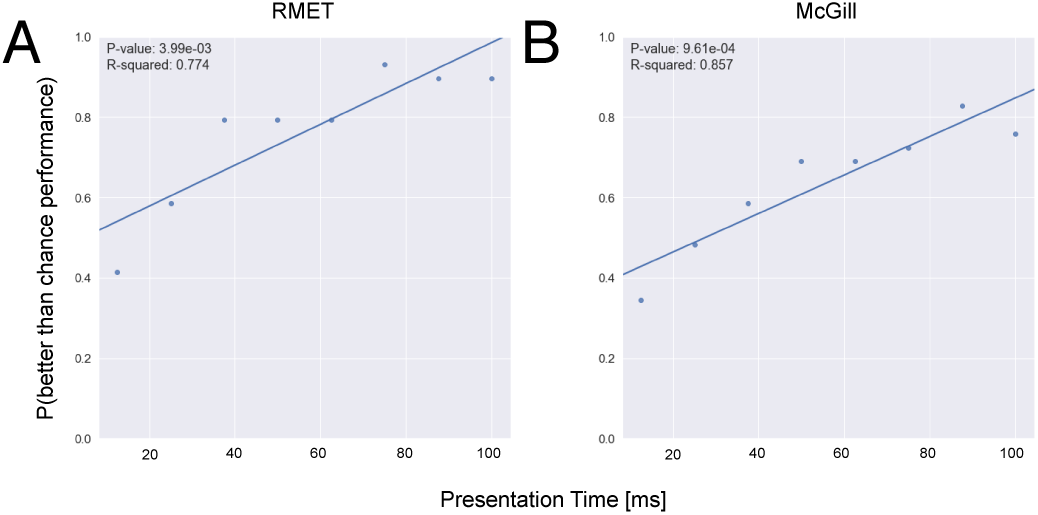
The figure shows the proportion of subjects performing better than chance for each presentation time for (A) the RMET and (B) the McGill Database stimuli.

The mean guess rate across subjects reported by the observers (see Table 1) is 50.4% (SD=18.5). Figure 3 compares individual estimations of guess rate in comparison to measured accuracy (RMET: Figure 3A, McGill Figure 3B). Linear regression tests found no correlation between these variables (*R*^2^ and *p*-values are shown in each graph in Figure 3). This suggests that participants are not able to make accurate judgements on their level of performance at these tasks.

**Figure 3.**
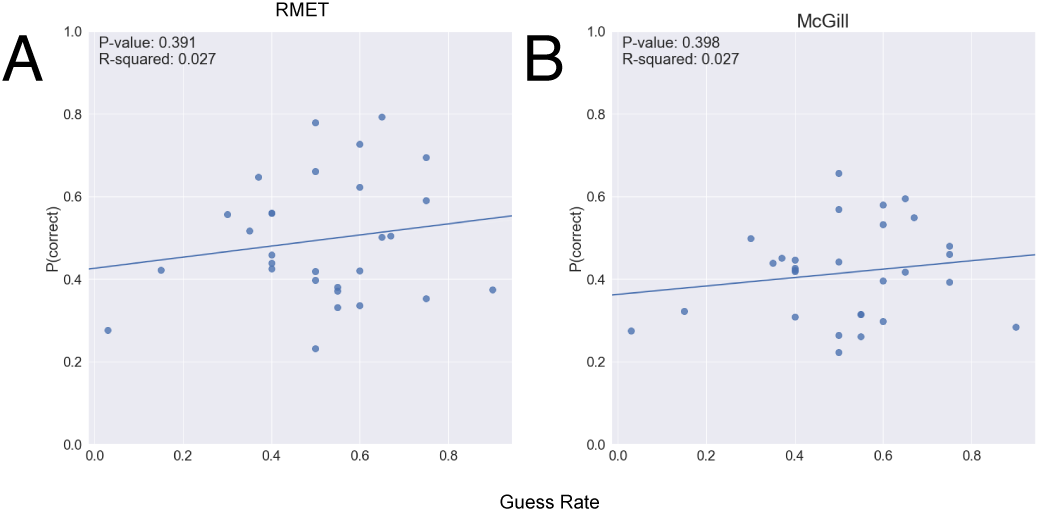
The figure shows response accuracy (proportion correct) as a function of guess rate for (A) RMET and (B) the McGill Database stimuli.

In an additional analysis, the target terms were grouped into positive and negative terms by consensus of the investigators. The following terms were considered to be positive: interested, playful, confident, desire, flirtatious, fantasizing, friendly; and the terms upset, worried, doubtful, accusing, nervous, suspicious, hostile, concerned, regretful, despondent, distrustful and uneasy were identified as negative. The remaining terms were excluded from this analysis because they were considered neutral or ambiguous (e.g., ‘defiant’ and ‘thoughtful’). Figure 4 shows the response accuracy as a function of presentation time for the RMET stimuli (Figure 4A) and the McGill stimuli (Figure 4B). Two-tailed *t*-tests show that for the RMET stimuli, the overall mean accuracy of negative terms and positive terms was statistically significantly different (*t*(2.450), *p* =. 044) (µ = 0.48,±0.08 *SD* vs µ = 0.50,±0.08 *SD*). For the McGill stimuli, this difference was larger (µ = 0.44,±0.08 *SD* vs µ = 0.37,±0.04 *SD*); *t*(−4.204), *p* =. 004).

**Figure 4.**
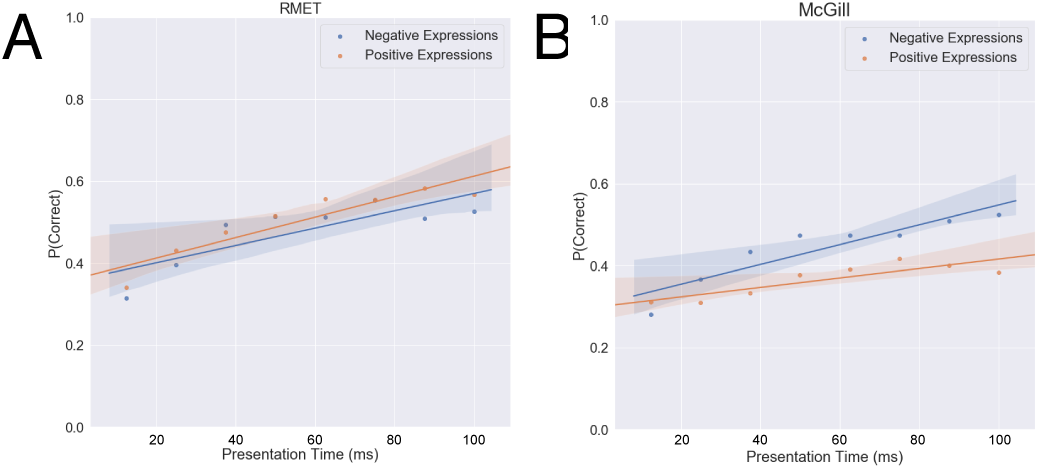
Negative vs Positive terms: The figure shows the response accuracy as a function of presentation time for the (A) RMET and (B) the McGill stimuli. The shaded regions represent 95% confidence intervals.

Figure 5 shows box-and-whisker plots of response times (considering only response times under 3s) for correct (orange) and incorrect decisions (blue) across all participants for the RMET (Figure 5A) and the McGill stimuli (Figure 5B). The middle line represents the median, the central box displays the interquartile range (IQR). The whiskers represent 1.5 times the IQR, and the points plotted outside of these whiskers representing the outliers. Table 3 and 4 show that for both stimuli classes, most response times were statistically significantly shorter for incorrect compared to correct decisions based on two-tailed *t*-tests for each target term. For the McGill stimuli, all mean response times were significantly different (*p* <. 05) in mean response time between correct and incorrect decision except for two emotions: confident and regretful. For the RMET stimuli, all mean response times were significantly different (*p* <. 05) except for two emotions: defiant and nervous. In an additional analysis, we analysed the effect size to investigate the magnitude in differences between correct and incorrect response times. The Cohen’s *d* of the observed response times for the RMET stimuli was 0.480 and 0.354 for the McGill stimuli. This shows that longer response times had a small magnitude effect (0.5 > *d* > 0.2) on increasing accuracy (Cohen, 2013).

**Table 3.** List of RMET target term response time two-tailed t-test results.

|  | **Target Term** | **Mean Response Time - Correct** | **Mean Response Time - Incorrect** | **Degrees of Freedom** | **t-statistic** | **p-value** |
| --- | --- | --- | --- | --- | --- | --- |
| 1 | defiant | 1.81 | 1.80 | 341 | -0.08 | 0.935639 |
| 2 | concerned | 2.05 | 1.69 | 335 | -5.11 | 0.000001 |
| 3 | reflective | 1.99 | 1.61 | 367 | -5.76 | 0.000000 |
| 4 | serious | 1.91 | 1.65 | 377 | -4.46 | 0.000011 |
| 5 | playful | 1.84 | 1.71 | 337 | -2.00 | 0.046495 |
| 6 | flirtatious | 1.92 | 1.69 | 370 | -3.58 | 0.000405 |
| 7 | friendly | 2.07 | 1.71 | 341 | -5.39 | 0.000000 |
| 8 | distrustful | 2.05 | 1.61 | 320 | -6.18 | 0.000000 |
| 9 | anticipating | 2.03 | 1.79 | 325 | -3.13 | 0.001948 |
| 10 | despondent | 2.00 | 1.62 | 337 | -5.67 | 0.000000 |
| 11 | thoughtful | 2.07 | 1.68 | 359 | -6.04 | 0.000000 |
| 12 | fantasizing | 2.03 | 1.63 | 633 | -8.23 | 0.000000 |
| 13 | suspicious | 1.99 | 1.69 | 366 | -4.84 | 0.000002 |
| 14 | upset | 1.80 | 1.61 | 396 | -3.44 | 0.000639 |
| 15 | confident | 2.05 | 1.57 | 329 | -7.16 | 0.000000 |
| 16 | desire | 2.03 | 1.71 | 328 | -4.46 | 0.000011 |
| 17 | regretful | 2.00 | 1.68 | 325 | -4.34 | 0.000019 |
| 18 | hostile | 1.79 | 1.68 | 374 | -1.80 | 0.072535 |
| 19 | worried | 1.92 | 1.65 | 358 | -4.25 | 0.000029 |
| 20 | pensive | 1.99 | 1.56 | 373 | -7.12 | 0.000000 |
| 21 | sceptical | 2.07 | 1.59 | 338 | -7.23 | 0.000000 |
| 22 | tentative | 1.98 | 1.76 | 292 | -2.91 | 0.003971 |
| 23 | uneasy | 2.06 | 1.66 | 337 | -5.76 | 0.000000 |
| 24 | insisting | 1.97 | 1.63 | 352 | -5.24 | 0.000000 |
| 25 | accusing | 2.01 | 1.60 | 323 | -5.86 | 0.000000 |
| 26 | preoccupied | 1.94 | 1.58 | 690 | -7.83 | 0.000000 |
| 27 | nervous | 1.74 | 1.72 | 349 | -0.25 | 0.802056 |
| 28 | doubtful | 1.98 | 1.76 | 349 | -3.18 | 0.001622 |
| 29 | decisive | 1.98 | 1.59 | 362 | -6.46 | 0.000000 |
| 30 | contemplative | 1.99 | 1.61 | 379 | -6.17 | 0.000000 |
| 31 | interested | 2.04 | 1.78 | 612 | -5.16 | 0.000000 |
| 32 | cautious | 2.01 | 1.71 | 658 | -6.10 | 0.000000 |

**Table 4.** List of McGill target term response time two-tailed t-test results.

|  | **Target Term** | **Mean Response Time - Correct** | **Mean Response Time - Incorrect** | **Degrees of Freedom** | **t-statistic** | **p-value** |
| --- | --- | --- | --- | --- | --- | --- |
| 1 | defiant | 1.85 | 1.68 | 377 | -2.40 | 0.017010 |
| 2 | concerned | 1.82 | 1.58 | 403 | -4.46 | 0.000011 |
| 3 | reflective | 1.86 | 1.62 | 381 | -3.74 | 0.000211 |
| 4 | serious | 1.91 | 1.53 | 402 | -6.18 | 0.000000 |
| 5 | playful | 1.77 | 1.62 | 390 | -2.40 | 0.016685 |
| 6 | flirtatious | 1.84 | 1.70 | 371 | -2.17 | 0.030735 |
| 7 | friendly | 1.81 | 1.67 | 390 | -2.26 | 0.024133 |
| 8 | distrustful | 1.88 | 1.59 | 372 | -4.75 | 0.000003 |
| 9 | anticipating | 1.87 | 1.66 | 368 | -3.26 | 0.001232 |
| 10 | despondent | 1.84 | 1.69 | 353 | -2.06 | 0.040587 |
| 11 | thoughtful | 1.88 | 1.60 | 350 | -4.32 | 0.000022 |
| 12 | fantasizing | 1.78 | 1.69 | 766 | -1.88 | 0.061096 |
| 13 | suspicious | 1.75 | 1.45 | 422 | -5.74 | 0.000000 |
| 14 | upset | 1.90 | 1.59 | 389 | -4.81 | 0.000002 |
| 15 | confident | 1.82 | 1.72 | 347 | -1.32 | 0.187118 |
| 16 | desire | 1.93 | 1.67 | 373 | -3.94 | 0.000105 |
| 17 | regretful | 1.71 | 1.66 | 356 | -0.69 | 0.488979 |
| 18 | hostile | 1.86 | 1.69 | 376 | -2.82 | 0.005139 |
| 19 | worried | 1.74 | 1.50 | 428 | -4.71 | 0.000004 |
| 20 | pensive | 1.86 | 1.52 | 389 | -5.76 | 0.000000 |
| 21 | sceptical | 1.95 | 1.57 | 382 | -6.46 | 0.000000 |
| 22 | tentative | 1.98 | 1.56 | 352 | -6.45 | 0.000000 |
| 23 | uneasy | 1.88 | 1.57 | 395 | -5.33 | 0.000000 |
| 24 | insisting | 1.90 | 1.64 | 381 | -4.25 | 0.000029 |
| 25 | accusing | 1.88 | 1.59 | 377 | -4.72 | 0.000003 |
| 26 | preoccupied | 1.83 | 1.58 | 722 | -5.01 | 0.000001 |
| 27 | nervous | 1.83 | 1.66 | 394 | -2.01 | 0.046405 |
| 28 | doubtful | 1.91 | 1.61 | 391 | -4.79 | 0.000002 |
| 29 | decisive | 1.78 | 1.65 | 373 | -2.08 | 0.038361 |
| 30 | contemplative | 1.81 | 1.59 | 407 | -3.85 | 0.000136 |
| 31 | interested | 1.85 | 1.68 | 707 | -3.20 | 0.001485 |
| 32 | cautious | 1.90 | 1.67 | 755 | -4.70 | 0.000003 |

**Figure 5.**
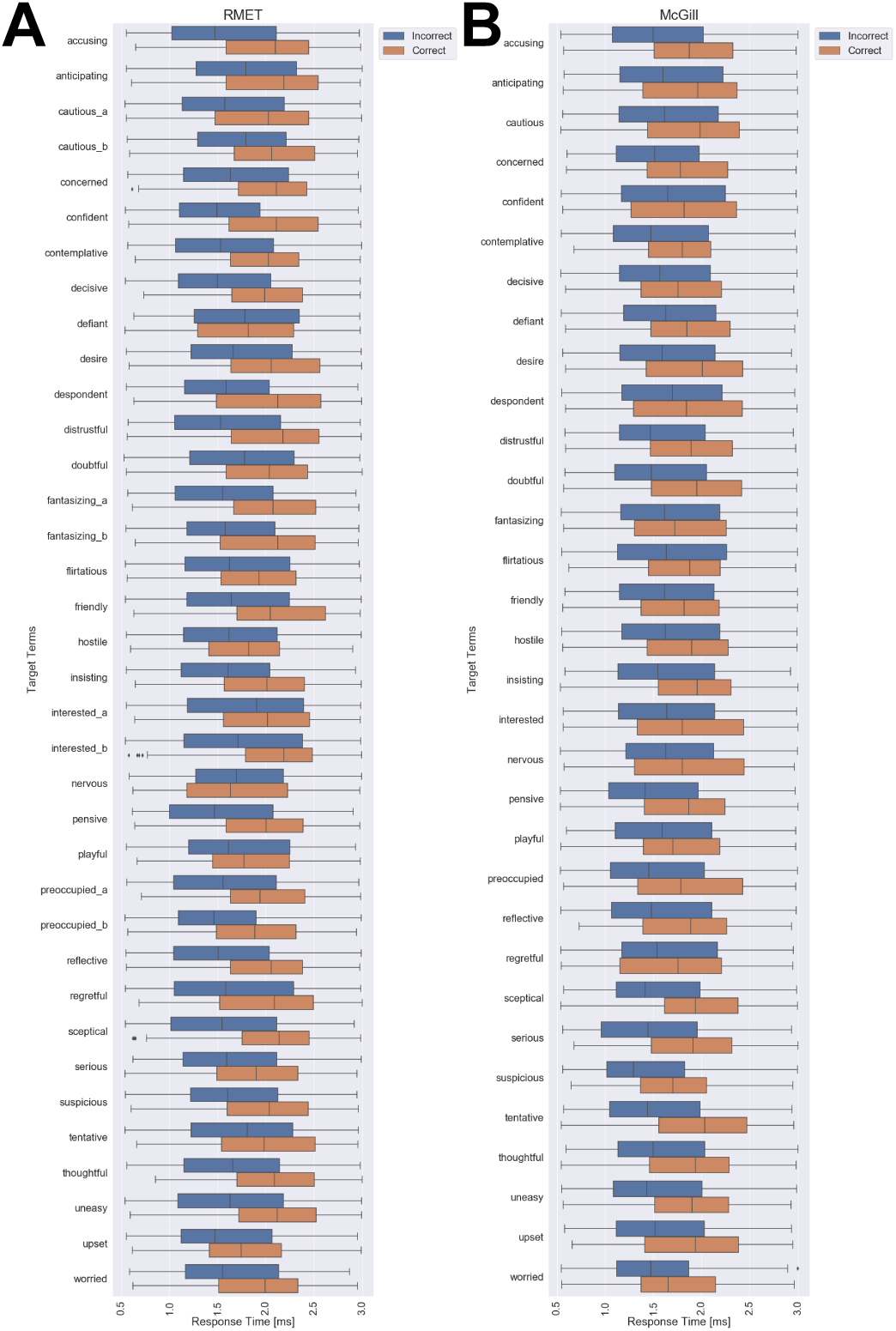
The figures show box-and-whisker plots of response times (considering only response times under 3 s) for correct (orange) and incorrect decisions (blue) across all participants for the RMET (A) and the McGill stimuli (B). The middle line represents the median, the central box displays the interquartile range (IQR), the whiskers being a function of 1.5 the IQR, and the points plotted outside of these whiskers representing the outliers.

In a final analysis, we show the relationship between correct and incorrect responses in a confusion matrix (RMET: Figure 6A, McGill: Figure 6B). The color-coding within this heat map represents the number of selections across all participants. The overall diagonal pattern in both figures shows that subjects frequently chose the correct term. Interestingly, the terms cautious, fantasizing, interested have been frequently identified, whereas preoccupied has been confused with the term puzzled for both stimulus classes.

**Figure 6.**
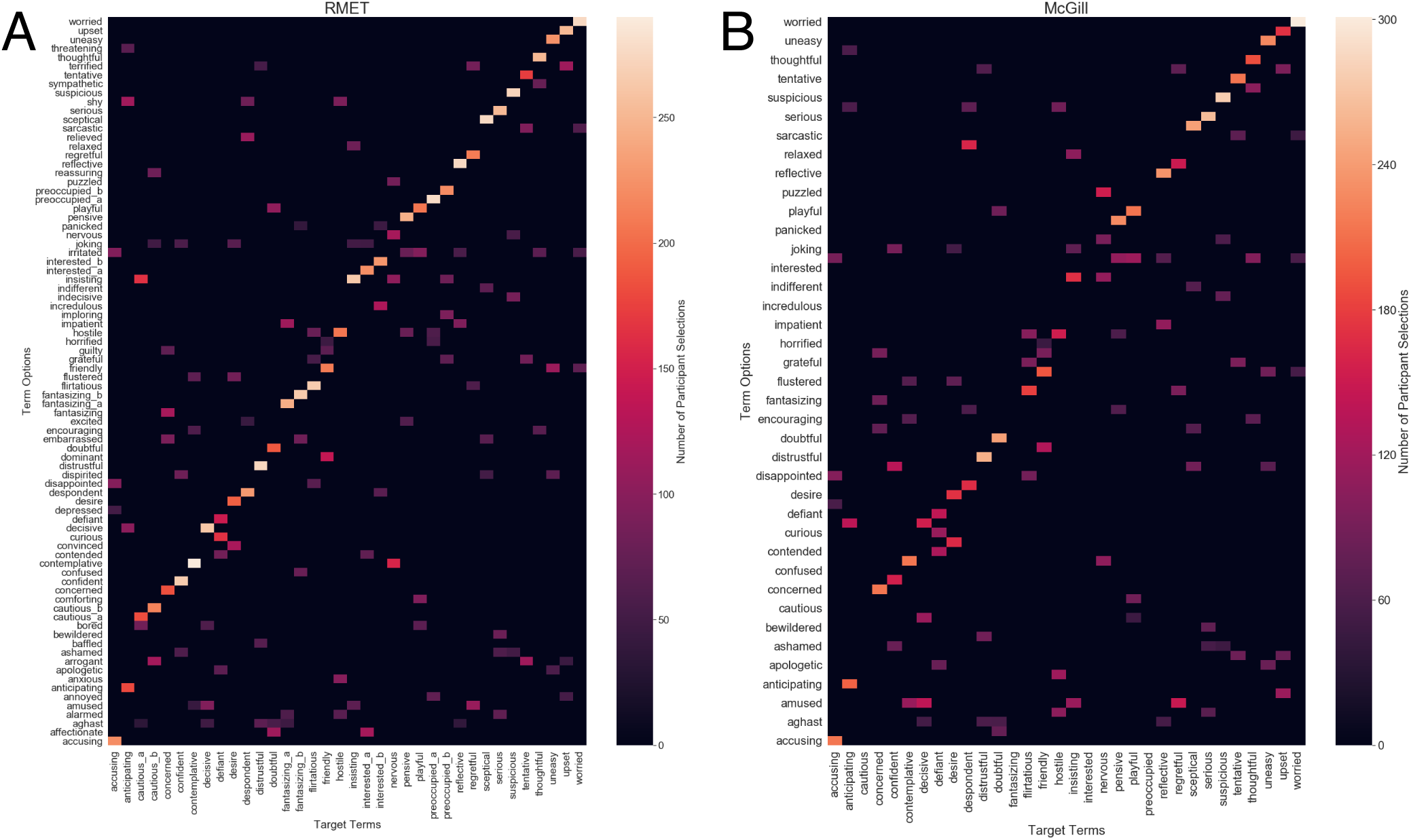
Confusion Matrix: (A) RMET and (B) the McGill Database stimuli. The color-code refers to the number of subjects.

## Discussion

It has long been acknowledged that there are a number of “low-level” components to Theory of Mind - for example, detection of direction of gaze and joint attention (see Baron-Cohen et al., 2001). Complex Theory of Mind, as modeled by Theory-Theory and Simulation-Theory, are cognitively demanding: they are computationally resource-intensive (Apperly, 2010) and would be expected to be slow and non-automatic. There are indeed many contexts in which theorizing about others’ mental states or putting oneself imaginatively in the position of another is both possible and useful; when the behavioural circumstances are unusual or complex - e.g., thinking about the motivations behind a politician’s campaign promises - it is typically necessary to be able to devote cognitive resources to reasoning about others’ mental states. In many contexts, however - landing a plane, passing the puck in hockey, improvising musically - one must be able to identify the mental states of others rapidly and automatically. The capacity of participants to perform better than chance on this task, even at presentation times of just 12.5ms, appears to be at odds with a construal of Theory of Mind as a slow and laborious process. Although Theory-Theory and Simulation-Theory may be useful models of some applications of Theory of Mind, therefore, they appear to be less well-suited to applications of the kind we report. This may point to a more substantive set of low-level components, beyond elements like direction of gaze, that play a role in Theory of Mind. The results of the current study may be better explained by a dual-process understanding of Theory of Mind. A number of investigators (notably Apperly and his colleagues; see Apperly, 2010, Apperly & Butterfill, 2009; see also, e.g. Leslie, 1994, Perner, 1991, Tager-Flusberg & Sullivan, 2000) have proposed that there are two distinct Theory of Mind systems or processes, subserving the ability to reason carefully about others’ mental states as well as the rapid, real-time capacity to represent those states in the course of social interaction, both of which are necessary for social cognition. So-called “dual process” or “dual systems” theories have proved useful in a number of areas of psychology (for overviews, see Evans, 2008, 2010, Evans & Frankish, 2009). The two processes or systems exhibit a cluster of (neither necessary nor sufficient) features: Type 1 cognition tends to be unconscious, implicit, automatic, effortless, fast, nonverbal, modular, domain-specific, and independent of general intelligence and working memory. In contrast, Type 2 cognition tends to be conscious, explicit, controlled, requiring effort, slow, non-modular, domain-general, and linked to general intelligence (Evans, 2008). The kind of cognition involved in successfully identifying expressions at very fast exposure times in this task is thus suggestive of a fast, automatic Type-1 cognition. The question of how Theory-Theory and Simulation-Theory might explain such cognition remains open. Our results show that there are no statistically significant differences between the RMET and the McGill stimuli with respect to accuracy of performance (see Figure 3). This is surprising, given that the RMET stimuli were selected because they were considered good representations of certain expressions, whereas the McGill stimuli were a result of actors being asked to “perform” certain expressions. Further, the RMET stimuli were selected for inclusion in a test which was specifically focused on the ocular region. The McGill stimuli, on the other hand, were captured to construct a test of facial expression discrimination in which all face features were visible. Accordingly, it could be proposed that, in general, the RMET stimuli contain more distinctive information within the ocular region than the McGill stimuli.

Here we should shortly reflect on the ecological validity of both approaches: whereas the more pronounced, more prototypical and in the end clearer stimuli from RMET might be more efficiently and more accurately processed, the McGill one might reflect more the typical conditions of everyday life requirements where people do not express the most prototypical but the most natural expression - and these are often much more subtle and partially executed than stereotypically acted and strongly selected stimuli. As noted above, the literature on which expressions are more salient - that is, which are more quickly and easily recognized - is mixed. Some have argued that positive expressions like happiness are more easily recognized, while others have argued that it is rather negative expressions like fear or anger that have greater salience (Calvo et al., 2014, Calvo & Lundqvist, 2008, Palermo & Coltheart, 2004, Tracy & Robins, 2008). Our results show no statistically significant difference in positive or negative expressions with respect to salience in the RMET stimuli. However, with the McGill stimuli, we found that performance for negative expressions improved much more rapidly than that for positive ones. This could be attributable to image-based aspects of the McGill stimuli. Specifically, Nummenmaa and Calvo (2015) proposed that contrast is a useful cue for rapid identification of expressions. If the McGill stimuli differ across positive and negative groupings in terms of contrast to a greater extent than the RMET, this could explain the differing results for positive and negative expressions with the two tests. Image analysis, however, revealed that this is not the case. Figure 7A shows Root Mean Square (RMS) Contrasts calculated for all RMET (red) and McGill (blue) stimuli. While there is significant variability in contrasts between various RMET stimuli, the McGill stimuli are uniform, with little difference between positive and negative expressions. Across the full range of stimuli (including both positive and negative) very little variation as can be seen in Figure 7A.

**Figure 7.**
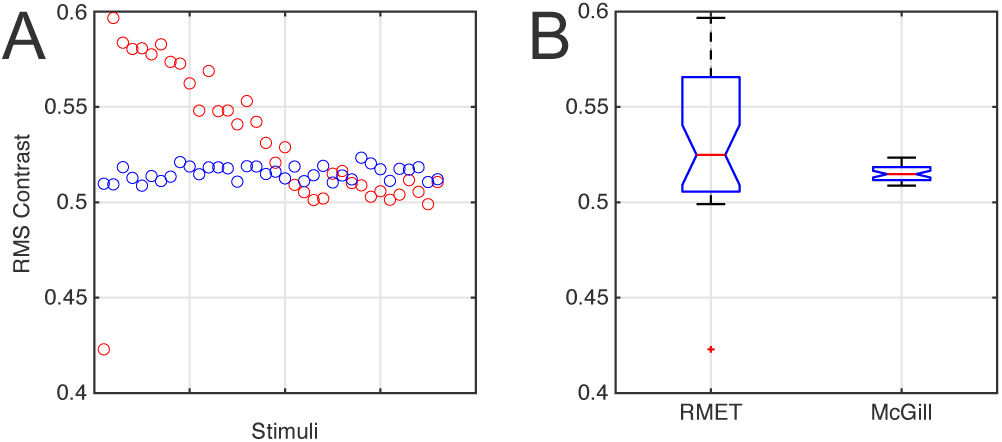
(A) shows Root Mean Square Contrast (RMS Contrast) calculations for all RMET (red) and McGill (blue) stimuli. (B) shows the corresponding boxplots for RMET (right) and McGill (left). On each box, the central mark indicates the median, and the bottom and top edges of the box indicate the 25th and 75th percentiles, respectively. The whiskers extend to the most extreme data points not considered outliers, and the outliers are plotted individually using the ‘+’ symbol.

It is noteworthy that face expression identification accuracy saturates on average around > 60% for both stimuli types (Figure 1). Restriction of available information to the eye region may partly explain this limitation of performance. It is well established that the eyes make a disproportionate contribution to the identification of emotional facial expressions (Baron-Cohen, 1997, Jack et al., 2012). Previous studies, however, have indicated that other face features (e.g. nose, mouth) also communicate information which facilitates interpretation of facial expressions (Baron-Cohen, 1997, Eisenbarth & Alpers, 2011, Yuki et al., 2007). This suggests that an improvement in accuracy may be achieved if the stimuli were adapted to include more face information.

## Acknowledgement

This research was supported by a Social Sciences and Humanities Research Council of Canada grant #435-2017-1215, 2017 given to I.G. and G.S.

